# Understanding biochemical design principles with ensembles of canonical non-linear models

**DOI:** 10.1101/2020.02.28.969170

**Authors:** Lukas Bromig, Andreas Kremling, Alberto Marin-Sanguino

**Affiliations:** Specialty Division for Systems Biotechnology, Technische Universität München, Boltzmannstraße 15, D-85748 Garching, Germany

## Abstract

Systems biology applies concepts from engineering in order to understand biological networks. If such an understanding was complete, biologists would be able to design *ad hoc* biochemical components tailored for different purposes, which is the goal of synthetic biology. Needless to say that we are far away from creating biological subsystems as intricate and precise as those found in nature, but mathematical models and high throughput techniques have brought us a long way in this direction. One of the difficulties that still needs to be overcome is finding the right values for model parameters and dealing with uncertainty, which is proving to be an extremely difficult task. In this work, we take advantage of ensemble modeling techniques, where a large number of models with different parameter values are formulated and then tested according to some performance criteria. By finding features shared by successful models, the role of different components and the synergies between them can be better understood. We will address some of the difficulties often faced by ensemble modeling approaches, such as the need to sample a space whose size grows exponentially with the number of parameters, and establishing useful selection criteria. Some methods will be shown to reduce the predictions from many models into a set of understandable “design principles” that can guide us to improve or manufacture a biochemical network. Our proposed framework formulates models within standard formalisms in order to integrate information from different sources and minimize the dimension of the parameter space. Additionally, the mathematical properties of the formalism enable a partition of the parameter space into independent subspaces. Each of these subspaces can be paired with a set of criteria that depend exclusively on it, thus allowing a separate sampling/screening in spaces of lower dimension. By applying tests in a strict order where computationally cheaper tests are applied first to each subspace and applying computationally expensive tests to the remaining subset thereafter, the use of resources is optimized and a larger number of models can be examined. This can be compared to a complex database query where the order of the requests can make a huge difference in the processing time. The method will be illustrated by analyzing a classical model of a metabolic pathway with end-product inhibition. Even for such a simple model, the method provides novel insight.

**Author summary:** A method is presented for the discovery of design principles, understood as recurrent solutions to evolutionary problems, in biochemical networks.

The method takes advantage of ensemble modeling techniques, where a large number of models with different parameter values are formulated and then tested according to some performance criteria. By finding features shared by successful models, a set of simple rules can be identified that enables us to formulate new models that are known to perform well, a priori. By formulating the models within the framework of Biochemical Systems Theory (BST) we manage to overcome some of the obstacles often faced by ensemble modeling. Further analysis of the selected modeling with standard machine learning techniques enables the formulation of simple rules – design principles – for building good performing networks. We illustrate the method with a well-known case study: the unbranched pathway with end-product inhibition. The method manages to identify the known features of this well-studied pathway while providing additional guidelines on how the pathway kinetics can be tuned to achieve a desired functionality – e.g. demand vs supply control – as well as to identifying important tradeoffs between performance, robustness and and stability.

## Introduction

Teleonomy is a concept coined by Norbert Wiener [1] and popularized by Jacques Monod [2] that refers to the appearance of design in biological systems. The tinkering of natural selection with biological systems on an evolutionary time-scale produces results that are very similar to what an engineer would design starting from first principles [3]. Following this philosophy, we can say that we will understand a biochemical network the day when we will be able to design it from scratch or modify it to fit our needs. Achieving this level of understanding is the goal of systems biology and being able to apply it is the long-term goal of synthetic biology.

In order to bridge the gap between description and understanding, mathematical modeling can be extremely useful. When a model is available, expensive and time consuming experiments can be replaced by simulations. An approach often used when modeling a biochemical network is to formulate a system of differential equations based on the mass balance:

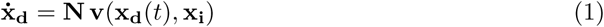

where **N**, the stoichiometric matrix, contains all the relevant information on the structure of the fluxes within the network. Vector **v** is a set of complicated non-linear functions describing growth rates, reaction kinetics, etc. The variables of this system collected in vector **x** are mostly metabolites but can include any kind of physicochemical entities such as temperature or volume. The vector is normally partitioned into two components: the dependent variables, **x**_**d**_, that will change according to the dynamics described by the equations, and the independent variables, **x**_**i**_, that will remain constant or have dynamics determined externally as forcing functions. In the context of this work, we will assume all independent variables are held constant.

Some of the parameters involved in Eq 1 are relatively easy to determine. Most coefficients of the stoichiometric matrix, for instance, can be inferred from the genome of an organism. Functional genomics enables the compilation of a list of the reactions taking place in the cell and in this way establish most of the entries of the stoichiometric matrix. Stoichiometric unknowns, such as the specificity of some reactions, can then be determined with uncomplicated experiments. Analytical methods based exclusively on the stoichiometric matrix – e.g. Flux Balance Analysis (FBA) — are extremely popular in spite of their many limitations. One of the reasons for this popularity is the very small amount of information needed to formulate these models. This ability of obtaining informative results from limited information comes from its use of optimality principles to define physiologically meaningful states. FBA explores every possible flux distribution within a network and finds optimal states – e.g. maximum biomass yield or maximum ATP production – these states are not only likely to be favored by evolution, but they are also immediately informative for the field specialist. Even if an organism does not fit into an optimal profile, it will very often reach a compromise between different optimal states. The linearity of FBA models makes it easy not only to compute their solutions, but also to combine them using superposition principles. Full dynamic models cannot benefit from any of these assumptions, the search of optimal states almost invariably results in NP-hard problems and the superposition principle does not apply. In spite of these difficulties, there is a long tradition of using dynamic models to determine biologically reasonable architectures of biochemical networks. This can be achieved by using structured modeling formalisms that simplify the mathematical treatment. Pioneering works from the late sixties and early seventies focused on discovering “design principles” [4] in biochemical pathways. A design principle is simply a problem solving strategy that can be reused in different contexts. Design principles lead to patterns that can be observed recurringly in nature, such as end-product inhibition, which is found very often in biosynthetic pathways and will be the object of our case study. Since these pioneering works relied on analytic methods, their application was limited to relatively simple models. These techniques were expanded in the early 2000’s to include statistical analyses of ensembles of models [5] so they could tackle higher degrees of complexity. Two decades later, a wide range of “design” principles have been discovered, explaining why certain patterns or regulatory signals keep emerging in completely unrelated networks [6] but the interpretation of complex models is still an unsolved challenge. The advances in machine learning experienced during this time and the availability of easy to use yet powerful tools to implement such methods opens up many new avenues to advance in this direction.

The task of elucidating design principles from ensembles of models poses three main challenges: 1) Ensuring that the generated models achieve a good coverage of the parameter space, 2) identifying good performing models – in our case, good biochemical “designs” – and 3) backtracking their good performance to a set of defining characteristics – e.g. parameter values. In this work, we will illustrate how the formulation of models adhering to a well defined mathematical formalism can simplify these tasks.

## Materials and methods

### Ensemble modeling

Formulating a full dynamic model poses a formidable challenge due to the numerous parameters involved in the rate laws – **v**(**x**_**d**_(*t*), **x**_**i**_) in Eq 1. These parameters cannot be measured simultaneously by any high-throughput technique and their individual determination *in vitro*, besides being extremely labor intensive, does not really inform about their *in vivo* values. Even though there are examples of high-quality large-scale models [7], these models tend to be one of many options that fit the available data equally well. In most cases, the data available about a certain process is compatible with a wide variety of alternative models. Despite promising developments that have been shown for small models [8], the uncertainty associated with the parameter values of a dynamic model tends to remain high due to its high complexity and identifiability problems. This challenge has inspired scientists to turn towards other approaches. One such approach is ensemble modeling, which copes with uncertainty by generating and analyzing multiple models [9]. Initially used to explore the effect of different initial conditions on the predictions of chaotic dynamical systems [10], the technique has generalized to many different fields like machine learning [11], where special emphasis has been put on how to combine the predictions of multiple models. By evaluating the performance of the different models under different criteria, a set of representative models can be selected.

The main difficulty faced by ensemble modeling is the high dimensionality of the parameter space. The total number of models that can be formulated for a given problem is often much larger than what can be realistically analyzed. For this reason, different acceleration strategies have been proposed such as testing the models for stability before fully analyzing them [12] or by replacing some parameters by otherwise accesible information. An example of the latter approach is anchoring the models to a flux distribution at the steady state [13, 14], which can be measured or calculated by means of FBA. By generating only models that admit that flux distribution as a steady state, the dimensionality of the parameter space is reduced considerably and the analysis focuses on more realistic models. In principle, the same could be done with the steady state metabolite concentrations, but some practical difficulties often prevent it. First, there are no established methods that predict a single set of metabolite concentrations from a theoretical perspective. Second, even when such concentrations are measured, the backwards calculation from concentrations to parameters is not always straightforward due to the non-linear nature of the rate functions. In the following sections, we will show the advantages granted by formulating the model using a specific (power-law) formalism: First, the steady state distribution of fluxes and metabolites can be set *a priori* [15]. Second, the parameter space can be partitioned into sub-spaces that can be tested independently. Third and final, the parameters of the power-law formalism are more directly related to systemic responses than traditional parameters such as saturation constants. This simplifies the stages of model selection and interpretation.

### The power-law formalism

The use of simplified rate laws can minimize the number of parameters of the system and provide structural regularity. Using these rate laws around a reference state results in good approximations for more complex expressions, such as traditional Michaelis-Menten kinetics (see [4] and supplementary information S1). A classic example of such expressions is the power-law:

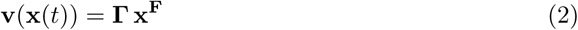

In this notation, 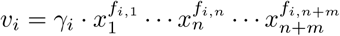, where *n* is the number of dependent and *m* that of independent variables. **Γ** is a diagonal matrix containing the positive constants *γ*_*i*_ that are called rate constants in analogy to the very similar Mass Action (MA) rate law. Also in analogy to MA, the parameters *f*_*i,j*_ are called kinetic orders but unlike the case of MA kinetics, the kinetic orders of a power-law are real numbers. For a more detailed discussion of this notation, see [16]. Systems described by a combination of Eq 1 and 2 are called Generalized Mass Action systems (GMA) [4, 17]. Rate constants can be set according to the formula

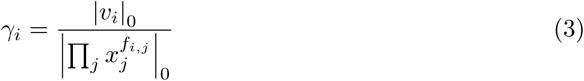

where the notation|·|_0_ represents a magnitude at a reference steady state. The importance of this definition is that the full dynamic behavior of the system around a certain steady state is fully specified by three sets of parameters: The flux distribution at the steady state, the corresponding vector of concentration profiles, and a set of kinetic orders **F**. A great advantage of this approach is that measuring flux distributions and metabolite concentrations *in vivo* is much easier than measuring kinetic parameters, so the above-mentioned strategy of “anchoring” models to a certain flux distribution and a certain profile of metabolite concentrations [18] happens naturally in this formalism. Using easy to measure values to reduce the dimensionality of the parameter space has proven to be a very effective strategy for ensemble modeling [19] that works particularly well with the power-law formalism [20]. Using models based on the power-law formalism or similar approaches such as Metabolic Control Analysis (MCA) offers a series of advantages [21] and does not result in a loss of generality. Once a set of the above-mentioned parameters is obtained, it can always be translated to more traditional biochemical parameters such as Michaelis constants or Hill coefficients [22, 23].

### Design space and criterion space

Introducing two concepts from optimization theory: the design space and criterion space will help to address the three challenges mentioned in the previous section, namely the formulation, selection, and interpretation of mathematical models.

The **design space** in an optimization problem is spanned by the variables that can be controlled and describes all feasible forms in which the model can be implemented. In our case, the parameters of a biological model reflect two important biological entities. On the one hand, parameters such as kinetic constants follow from protein sequences, cell architecture, etc. so they can be considered a proxy for the genotype. On the other hand, the rest of the parameters reflect constraints on the system, such as availability of a nutrient, equilibrium constants, etc. These parameters reflect the environment or more generally the ecological niche of the system. Together, the design space of the model defines all possible genetic variations of the system under a variety of environments [24, 25]. The idea of a design space follows naturally from the engineering analogy that motivates systems biology and played an important role in some of its foundational works [26]. In fact, this idea has established itself to the point of permeating into popular culture as can be seen in the following excerpt from a science fiction novel: “Similar pressures yielded similar solutions, just the way flight had evolved five different times on Earth. *Good moves in design space*” [27].

Just as the design space contains the relevant information about how a system is built out of its components, a counterpart is needed to describe the system as a whole and its interactions with the environment. This is done in engineering and optimization problems by quantifying its performance according to certain criteria. The set of scores for each case is then a point in the **criterion space**. In our case, the criteria will be evolutionary relevant goals such as the cost of maintaining a flux or the responsiveness of a pathway. In the simplest case, only one criterion is used and all systems can be ordered from best to worst performance. For more than one criterion, as is usually the case [28], there is a correspondence between each point of the design space and a point in the criterion space that describes its performance. Going back to the biological case, this correspondence reflects how each genotype in a certain environment (design space) produces a different phenotype (criterion space) [24].

Thus, the three tasks enumerated at the begining of this section can be reformulated as: sampling points in design space, selecting their images from criterion space, and find rules to trace these optimal phenotypes back to design space. Each of these three tasks will be explained in detail in the three subsequent sections.

### Sampling the design space

The number of samples needed to cover an n-dimensional parameter space with a certain density, *d*, grows exponentially with the number of parameters. The selection step that follows the sampling normally requires all parameter values simultaneously, thus leading to a bottleneck where a computationally expensive operation – e.g. numeric integration – is applied to a large number of parameter vectors. Sequential filtering of samples can simplify this task. For instance, a pre-selection of models with stable steady states before performing numerical integration can reduce the amount of operations significantly [12, 29]. This strategy can be taken one step further by dividing the design space into subspaces and sampling them independently. A system with two subsets of parameters of dimensions *p* and *q* will no longer require *d*^*p*+*q*^ samples to be covered, but rather *d*^*p*^ + *d*^*q*^. This reduces the number of necessary tests by several orders of magnitude and each individual test will be faster since it uses a restricted set of parameters. By formulating the model using the power-law formalism, models can be tested for a number of criteria using only a subset of the parameters of the system, thus enabling such a partition of the design space. The transition times of the metabolites, for instance, only depend on the flux distributions and the metabolite concentrations. The thermodynamic efficiency of an enzyme only depends on its proximity to equilibrium, which in turn is a function of the concentrations of its metabolites. Sensitivities, on the other hand, depend exclusively on kinetic orders and these are independent of the fluxes and metabolite concentrations as long as the reactions are far from equilibrium.

So, instead of generating a single ensemble of models where each instance has one dimension per parameter, three independent ensembles can be created. These ensembles, which we will call v-, x- and k-ensembles will contain flux distributions, metabolite concentrations, and kinetic orders, respectively. This partition of the design space allows a much denser sampling of each of the subspaces, from which models will be selected according to specific criteria for each ensemble. Only after filtering by these simple criteria, the selected samples from each subspace can then be combined to provide a much reduced set of full systems that can be subject to more computationally expensive tests. This process is akin to a complex database search composed of several criteria. If the most restrictive criterion is applied first, searches for all subsequent criteria will have to deal with fewer records and the whole process will be significantly faster. By analogy, this method achieves computational efficiency by formulating the final ensemble of models as the cartesian product of three sub-sets (v-, x- and k-ensembles) and filtering each subset separately.

There are many algorithms available to sample fluxes, concentrations and kinetic parameters in general. Although any of them are, in principle, compatible with the method presented here, some guidelines should be followed in order to guarantee ensembles that are consistent and separable. Flux distributions for the v-ensemble should not be drawn from purely stoichiometric approaches, since that may lead to thermodynamic inconsistencies. There are methods to obtain thermodynamically sound flux distributions [30] that have the advantage of also providing the necessary thermodynamic information to evaluate the x-ensemble or generate the ensemble itself [29]. Excessive proximity to equilibrium by any reaction would affect the kinetic orders and prevent an independent sampling of the k-ensemble. Fortunately, dealing with these cases is straightforward and can be done automatically by thermodynamic shortening [31]. Taking this into account, every kinetic parameter can be sampled easily from a uniform distribution with clearly defined limits. For instance, substrates and products usually have kinetic orders in the ranges (0,1) and (−1,0) respectively. Allosteric effectors can be as high in magnitude as their respective Hill coefficients.

### Model selection in criterion space

#### Selection criteria

There is a wide agreement among the systems biology community that some criteria have to be fulfilled by biological systems to be viable [5, 32, 33]. Many of these criteria can be computed using only one or two of the three ensembles discussed above, thus enabling a search for fit systems confined to a sub-space of both the design and criterion spaces.

- Goals depending on the v-ensemble Given a stoichiometric matrix, there is a variety of methods that calculate optimal flux distribution with respect to rates and yields. An example of such techniques is FBA, which has often been used to find optimum growth rates or biomass yields. Due to the enormous success of these techniques, constraint-based models can be considered a field in itself and a wide range of tools are available. Exploration of the v-ensemble, be it through optimization or sampling, has been thoroughly covered elsewhere [32], and it is normally done prior to further analysis. We will therefore focus on the other two ensembles and assume that a starting flux distribution or a discrete number of them have already been selected.
- Goals depending on the x-ensemble
  – Total concentration of metabolites. The total concentration of solutes in the cytoplasm is limited by their solubility limit as well as their toxicity. This places a hard upper limit on the possible values that each of the individual concentrations can reach [34]. Since overall solute concentration is a limited resource, the sum of all metabolites of an efficient pathway would be expected to be a goal for minimization and thus as low as possible.
  – Thermodynamic efficiency of the enzymes. Practically all enzyme rates can be factored [23, 33] as a constant term multiplied by a series of factors in the form:

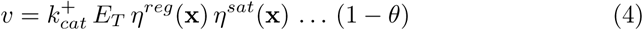 The maximum reaction rate of the reaction is the product of the catalytic constant *k*_*cat*_ times the amount of enzyme *E*_*T*_. All the other factors are bound between 0 and 1. Each of the *η*^·^(*x*) functions accounts for a different effect of the metabolites on the reaction rate – e.g. allosteric inhibition, saturation, etc. – and depends also on a series of parameters like Hill coefficients, saturation constants, etc. Finally, *θ* is the distance to equilibrium and will be bound between 0 and 1 for a reaction progressing left to right. That means that the thermodynamic term in the parenthesis determines the percentage of enzyme that will actually contribute to the net reaction, so it is often used as an orientative indication of the overall thermodynamic cost [33]. We will aggregate these terms in the criterion:

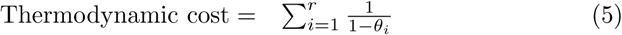

where *r* is the number of reactions to classify models according to thermodynamics. This criterion can be easily adapted to include information about catalytic efficiency for different steps in the pathway. See [31] for a discussion on how to use this type of kinetics together with power-laws.
  – Ratio of final products to intermediates (draft) Many intermediary metabolites are just necessary chemical steps to reach a product. Given the above-mentioned upper bound to the total content of the cell, a high concentration of any intermediate will lower the maximum tolerable concentration of the end product. Even though it is difficult to determine which metabolites are “useful” for the cell, some rules of thumb can easily be found. For every unbranched sequence of reactions in a metabolic pathway, only the final product may provide an advantage by having a high concentration, be it due to mass action or by providing exergonism for subsequent branches. The cell has much to gain from keeping the concentrations of all the “internal” metabolites as low as possible, since even a potential role in a regulatory interaction out of the branch can be performed at low concentrations. Thus, the ratio of internal intermediates to final product will be a measure of how much of the solubility limit of the cell is occupied by non-critical compounds. In order to assign a name to this criterion, we will use a nautical metaphor. The draft of a ship defines the depth reached by the fraction of its hull that is under water, so we will define

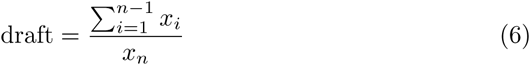

as the ratio of metabolites “submerged” inside a linear sequence of reactions to its final product. Draft minimization is therefore a worthy goal to be pursued in the design of a pathway. A straightforward generalization for more complex models would be to assign a draft to each linear segment. These segments can be identified from the stoichiometric matrix by calculating the metabolite adjacency matrix and removing all metabolites with connectivity greater than two. This will leave linear segments as connected components of the corresponding graph. In addition to these, an additional goal can be defined that depends on the combination of v and x-ensembles.
- Goals depending on the x- and v-ensemble
  – Transition time. The ratio of accumulated mass to throughput is often called transition time and it provides a reference for the time scale of the pathway [35].

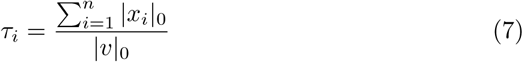
- Goals depending on the k-ensemble The sensitivities of the system depend almost exclusively on the kinetic orders – see Supplementary Information–, and as long as the reactions are far away from equilibrium, kinetic orders can be considered to be independent from the other parameters. These assumptions are not restrictive since it has been already shown that reactions close to equilibrium can be automatically eliminated from the sensitivity analysis through thermodynamic shortening of the pathway [31]. There is a long standing framework for sensitivity analysis of power-law systems, that is summarized in the supplementary information, based on the definitions of sensitivities (*S*) and log-gains *L*.

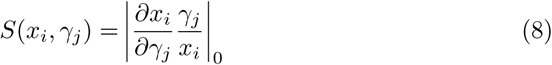

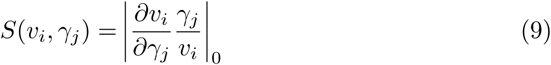

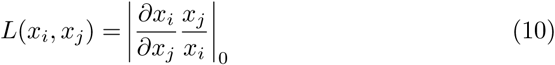

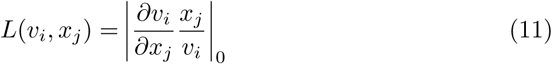 Sensitivities, *S*(*x*_*i*_, *γ*_*j*_) and *S*(*v*_*i*_, *γ*_*j*_) are the responses of the steady state of the system (dependent variables or fluxes) to perturbations on its parameters while logarithmic gains, *L*(*x*_*i*_, *x*_*j*_) and *L*(*v*_*i*_, *x*_*j*_), define the corresponding responses to changes in the independent variables. It is normally the case that a system has to be robust against changes in the parameters and therefore have low sensitivity, while it may be desirable that the system has a strong response to certain stimuli, so in many cases high gains are desired. The sensitivities and logarithmic gains of a system are determined by the kinetic orders and are, in general, independent of metabolite concentrations [4, 17], which enables separate sampling of the k-ensemble and selection based on sensitivities. The sole exception to this rule is the presence of moiety conservations. Metabolites involved in a moiety cannot change independently and the fraction of the moiety that is contained in each of its components – e.g 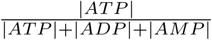 – has an effect on the sensitivity matrices. Dealing with conserved moieties does not pose an obstacle for the method since the number of metabolites involved in such moieties is a very small fraction of the total, specially for large networks. The matter can be dealt with by adding one degree of freedom to the k-ensemble in the form of a dummy variable for each such metabolite – e.g. ATP, ADP,… — these dummy variables will take values between 0 and 1 to represent the fraction of the moiety in that particular form. The dummy variables enable the calculation of the full sensitivity set for the system and establish a common key between the k- and the x- ensembles while allowing the separate sampling of both. A detailed account of sensitivity analysis is included in the supplementary information for the convenience of the reader.
- Goals that depend on all ensembles Stability of the steady state is a key necessity of any biochemical system and is often used as main criterion for the selection of biologically feasible models ([12]). Other criteria can be determined from numeric simulations of the system. Determining dynamic behavior of a system or the stability of its steady state requires the use of all parameters simultaneously and computationally expensive operations, so it is highly recommended to use other criteria to filter a large amount of models before evaluating this set.

#### Selection strategies

Performance indicators such as those defined above, can be used to discover “good” biochemical designs. As long as the criterion space has a dimension higher than one, a multiplicity of selection strategies can be followed.

1. Behavioral classes Alves et al. have proposed to categorize pathways according to their structure and behavior [5]. Establishing a cutoff value for a certain goal, a class can be defined by all the systems that perform above this value. This type of filtering can be applied to more than one criterion at a time, so optimal models can be defined as the behavioral class that performs above certain thresholds for a set of relevant criteria.
2. Pareto front Alternatively, the concept of Pareto optimality can be used for selection. A solution is efficient or Pareto optimal if there is no other solution that performs better for all criteria. The set of all efficient solutions, or Pareto front, collects all possible tradeoffs between different criteria. We will sometimes approximate the Pareto front by choosing all the efficient solutions in our sample by direct comparison. Occasionally, we will introduce a tolerance *ϵ* in such comparison, admitting solutions that are close enough to the Pareto front. The set of solutions so obtained will be named *ϵ*-Pareto front. In order to better characterize the Pareto front, we will sometimes resample by calculating convex combinations of efficient solutions. Such solutions will often be at or very close to the Pareto front, thus enriching our sample. The combination of epsilon selection and enrichment enables to obtain a virtually unlimited number of points on or very close to the Pareto front.

These two approaches can be used together. In this work, we will start most model selections by filtering out the least robust 50% element of every set, since robustness is a well-known requisite for the operation of biological systems [36, 37]. The remaining solutions will then be tested for Pareto optimality. The use of Pareto fronts to analyze biological performance has been very successful in the past [38], especially in cases where there is a monotonic relationship between genetic traits and their respective phenotypes [39].

### Tracking optimal phenotypes back to design space

Once a set of good designs has been identified, their optimality has to be explained in terms of their composition. In other words, their location in criterion space has to be traced back to some pattern in design space. Many machine learning techniques can be used for this purpose, here we will combine two of the simplest and best known tools for classification and dimensionality reduction.

#### Logistic regression

For each defined behavioral class, a logistic classifier was trained to identify its members by their position in design space. For each criterion, the algorithm – and similar ones such as Support Vector Machines – defines a hyperplane in design space separating it in two halves, one containing a majority of members of the class and one containing a majority of non-members. Although this separation is far from perfect, the vector normal to the hyperplane provides an optimal direction in the design space to separate alternative designs according to their parameter values. The normalized vector for a given criterion assigns a coefficient to each parameter. Parameters that tend to be high for class members will be assigned a positive coefficient and those that tend to be low, a negative one. The magnitude of such coefficient is also important with influential parameters (for better or for worse) having coefficients that are high in absolute value. Parameters that have a negligible influence on performance, however, will have coefficients with magnitudes well below the average. The python machine learning toolbox *scikit-learn* was used as described in the supplementary information. The reader is referred to [40, 41] for more information on *scikit-learn* and its implementation.

#### Principal component analysis

PCA is a very common technique for dimensionality reduction and has become a standard method in the fields of bioinformatics and biosystems engineering [42]. When the Pareto front is viewed in the design space, it is difficult to deduce a pattern or correlation. Thus, PCA is performed on the Pareto-efficient datasets using the above-mentioned sklearn toolkit. PCA provides a new set of coordinates in the design space that partition the variance in uncorrelated principal components (PCs), each with a weight associated to it. The contributions of the different parameters to each PC can be interpreted like those in logistic regression with parameters contributing to the high variance PCs being involved in differences between members of the Pareto front and parameters contributing mostly to low variant PCs being relatively constant. There are two main benefits in this procedure: Interactions between parameters and their respective weight become apparent and, if applicable, the dimensionality of the problem at hand can be reduced by rejecting insignificant principal components.

The overall procedure introduced is summarized in Fig. 1. The benefits of partitioning the parameter space become clear. The overall workflow is non-linear and can be adapted or reconfigured. Random resampling or generation of new datasets on the basis of PCA or convex combinations is possible along every step of the procedure.

**Fig 1.**
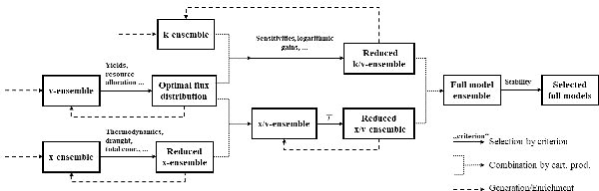
Summary of the workflow. A schematic view of the various steps towards a large-scale pathway analysis with the proposed ensemble modeling approach. The individual ensembles (Boxes with x-, k-, v-ensemble) are generated (--) and tested(—) separately. A reduced set of its members is then selected for further analysis. Mixed ensembles (Boxes with x/k-, v/x-ensemble) are obtained by generating all possible combinations (····) of the selected members of the former ensemble partitions. At the end of the process, the amount of models actually processed is the cartesian product of the small subsets of selected models in each ensemble. This subset is but a small fraction of all the possible combinations that have thus been proven inadequate at an early stage without the need for extensive testing or excessive computationally expensive operations.

### Case study

The concepts introduced in this work will be illustrated with a well-studied example, an unbranched pathway with end product inhibition [5, 31, 43, 44]. This type of pathway, shown in Fig. 2, is ubiquitous in biochemical systems and the reasons for its prevalence have been studied for more than forty years. After in-depth analyses using analytical methods [4, 26, 45–47], the unbranched pathway became a standard benchmark, and has since been used to exemplify a series of ensemble modeling approaches [22, 43, 48]. Simulations were done with pathways of different lengths but the results were extremely similar, so the method will be shown with a three-metabolite pathway for the sake of clarity. Thus, dependent variables *x*_1_ and *x*_2_ will be intermediates, the final product will be *x*_3_. The model has two independent variables: the concentration of precursor *x*_0_ and a dummy variable *x*_4_ introduced to reflect changes in cellular demand for product (*f*_4,4_ = 1).

**Fig 2.**
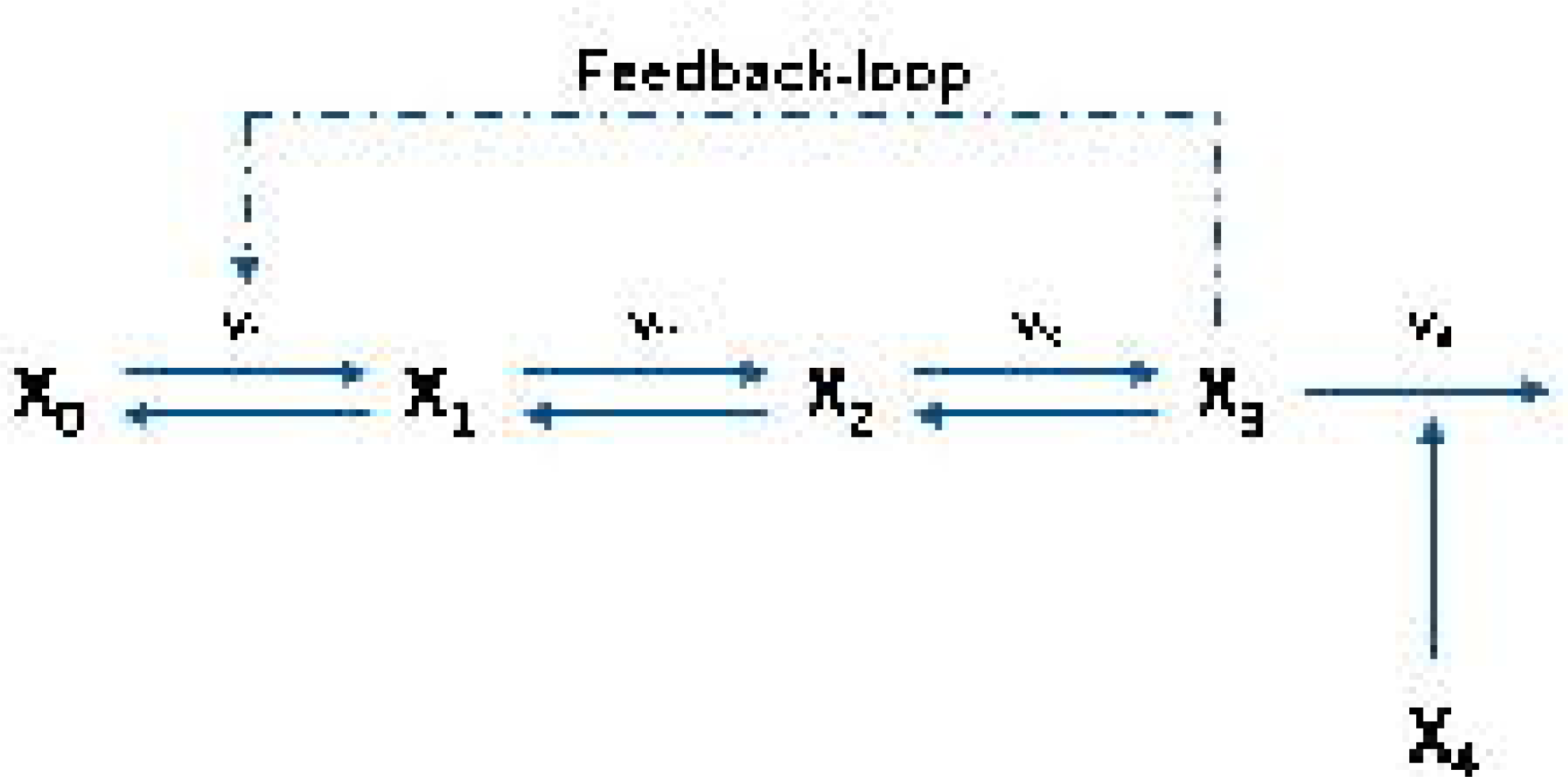
The model pathway. A three-step unbranched pathway with negative feedback loop is chosen as the model pathway of this case study. The selected pathway, with the initial precursor concentration *x*_0_ and the final product concentration *x*_3_, is reversible at every intermediary step. The dependent variable concentrations of the metabolites are named *x*_1_ to *x*_3_ and an additional variable *x*_4_ has been added to reflect variations in product demand.

#### Design space

The design space for this system has 12 dimensions and it can be partitioned into a one-dimensional v-ensemble – all reaction rates must be equal to the flux through the pathway —, three-dimensional x-ensemble: *x*_1_, *x*_2_ and *x*_3_, and an eight-dimensional k-ensemble: *f*_1,1_, *f*_1,3_, *f*_1,0_, *f*_2,1_, *f*_2,2_, *f*_3,2_, *f*_2,3_ and *f*_4,3_.

#### Criterion space

The system will be evaluated using the criteria described above. For the case of thermodynamic efficiency, equilibrium constants of all reactions will be assumed to be 1. In the case of sensitivity analysis, seven criteria were selected.

Three criteria reflect the robustness of the pathway: Aggregate Parameter Sensitivity (APS), Intermediates Sensitivity to Demand (ISD) and Intermediates Sensitivity to Supply (ISS).

1. APS. An overall measure of the robustness of the system.

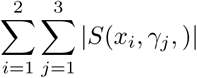 APS reflects the accumulation or depletion of the intermediates, *x*_1_ and *x*_2_, in response to fluctuations in enzyme activity.
2. ISD. The effect that a change in demand will have on the concentrations of intermediates.

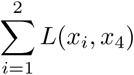
3. ISS. The effect that a change in supply (concentration of precursor *x*_0_) will have on the concentrations of intermediates.

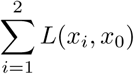 The four remaining criteria reflect the responsiveness of the pathway. Of these, two criteria characterize the pathway response to an increase in supply: Flux Supply Gain (FSG) and Product Supply Gain (PSG) with their counterparts reflecting the response to an increase in demand: Flux Demand Gain (FDG) and Product Demand Gain (PDG).
4. PDG. The impact that a change in product demand will have on its concentration.

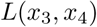
5. FDG. The flux response to an increase in demand.

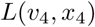
6. PSG. The increase in final product following an increase in precursor concentration:

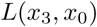
7. FSG. The flux response to an increase in precursor concentration.

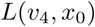

## Results

The workflow was applied to the unbranched pathway model using an arbitrary value of 31.5 mM/min for all the fluxes in steady state. For the model pathway under investigation, a reasonable flux through a biosynthetic pathway of a growing bacteria with a duplication time of *t*_*d*_ = 20 *min* was chosen. By estimating a reasonable flux, it is ensured that one stays within realistic bounds regarding the presented example. Further details regarding cellular composition and amino acid abundance are provided in the supplementary information [49].

The values for steady state metabolite concentrations were uniformly sampled in the interval 0.1*µM* − 10*mM*, which is a realistic cytosol concentration range. The values of the kinetic orders were uniformly sampled in the intervals [-1,0] for the backwards reaction (*f*_1,1_, *f*_2,2_, *f*_3,3_) and [0,1] for the forward reactions (*f*_1,4_, *f*_2,1_, *f*_3,2_, *f*_4,3_). Kinetic values between [-1,1] were chosen to represent a very basic enzymatic reaction; one without allosteric effects or multiple active sites. The kinetic order of the feedback strength *f*_1,3_ was sampled in the interval [-7,0]. Even though it is unrealistic to expect *in vivo* values below −2, large values for this kinetic order generate analogous results to simulating a longer pathway with realistic coefficients. For the sake of simplicity, this interval will be used instead of presenting the results of simulating pathways of different length, which yielded equivalent results. The demand factor *f*_4,5_ was fixed at 1, since it is part of a linear term. Unless otherwise specified, the equilibrium constants of the first three reactions will be considered to be one.

### Peculiarities of the x-ensemble

Unlike other parameters, the metabolite concentrations are subject to *a priori* thermodynamic constraints. For the pathway to be able to carry a flux, the ratios of products to substrates for all reactions have to be greater than the equilibrium constants. This results in the x-ensemble and the criteria based on it being biased even before a selection by performance is made. Fig. 3 shows the distributions of metabolite concentrations for the case study after filtering by feasibility for three different thermodynamic backgrounds. As can be seen, the distribution of feasible concentrations at the steady state can change dramatically depending on the specific thermodynamics of the pathway. This is of critical importance, since many modeling studies do not take thermodynamics into account prior to parametrizing their models.

**Fig 3.**
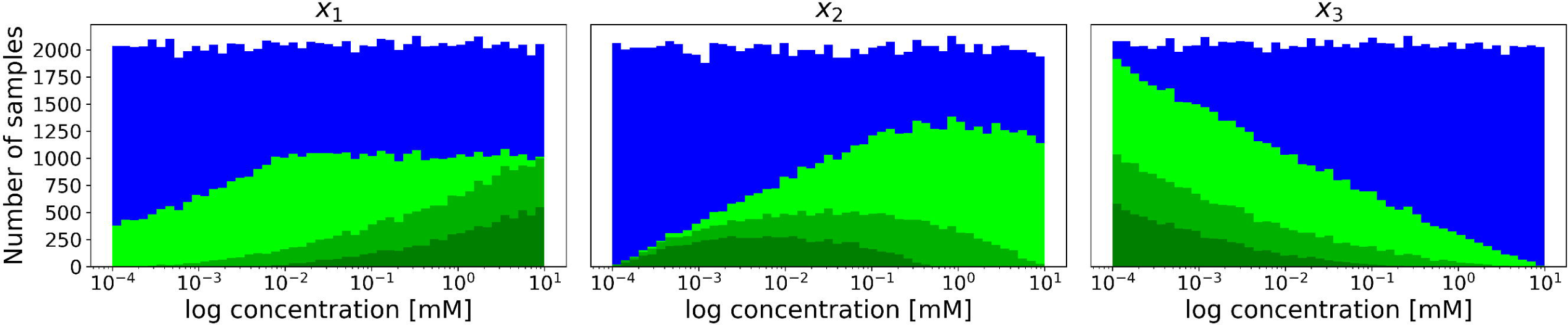
Thermodynamic feasibility analysis of the x-ensemble. The distributions of metabolite concentrations generated using a uniform distribution (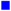) and the subset filtered by thermodynamic feasibility for different values of the second equilibrium constant: *K*_2_ = 1000 (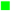), *K*_2_ = 1 (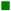), *K*_2_ = 0.05 (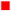). All other equilibrium constants are fixed at 1.

### Model selection in criterion space

In any multi-criteria optimization problem, the different criteria or goals will be conflicting at some point, but such conflicts can have different intensities or manifest themselves at different stages. While some goals are aligned to a high extent, others are clearly incompatible. Fig. 4, for instance, shows a sample of pathway designs represented as projections in two-dimensional criterion spaces. The criteria on the left panel ISD vs. FDG show a clear tradeoff where increasing the pathway response to demand involves a proportional increase of the sensitivity of intermediates. The right panel, however, shows that the two robustness criteria APS and ISS can both be brought almost to their optimal points simultaneously with some solutions being extremely close to the utopian point, where all goals achieve their absolute maximum.

**Fig 4.**
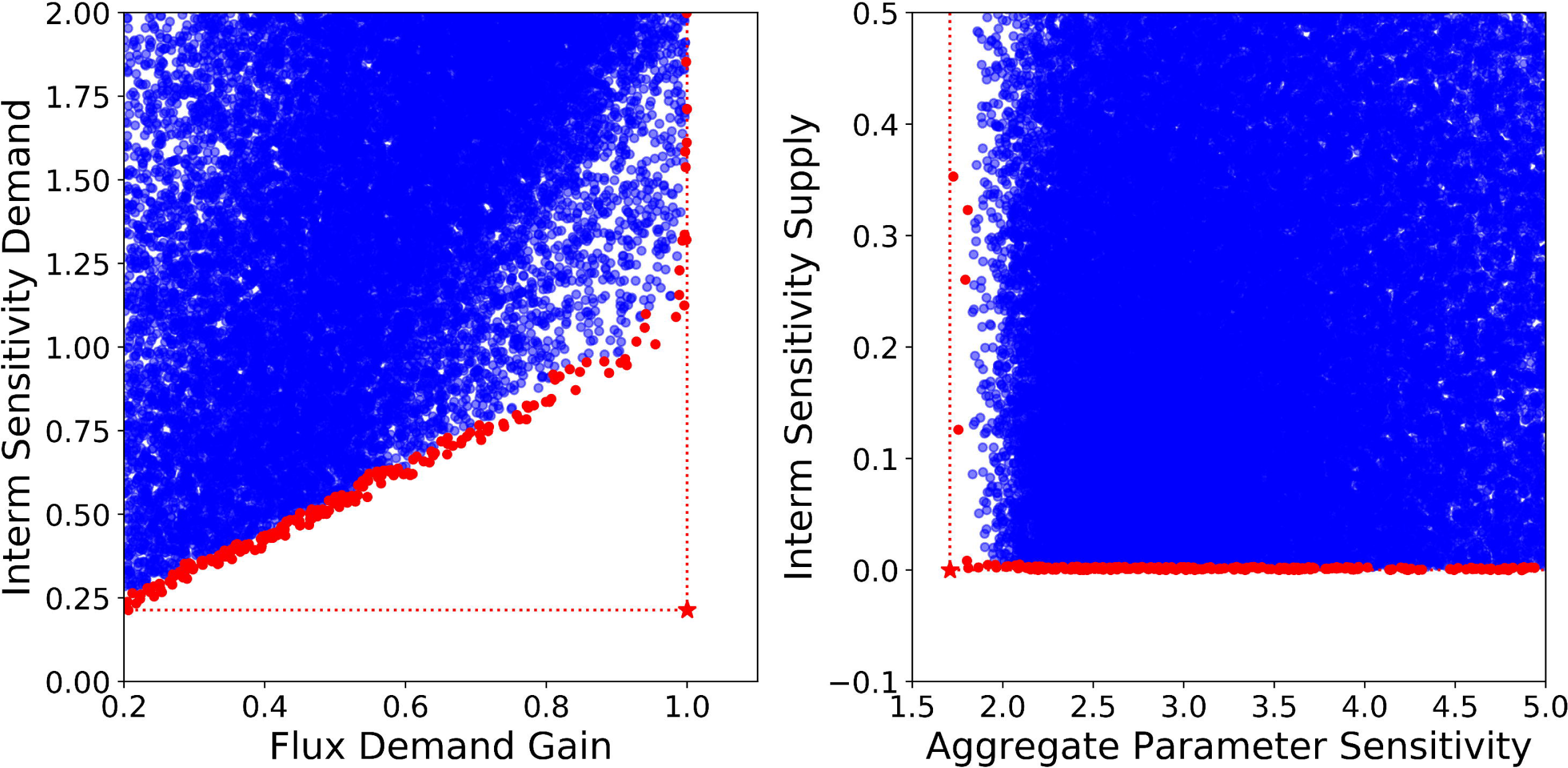
Examples of projections of the criterion space. Each model of the ensemble is represented by a point. The Pareto fronts are calculated for selected goal combinations. Depending on the number of selected goals, 2-dimensional or multi-dimensional Pareto frontiers can be calculated. The datasets are then grouped into the non-efficient datasets 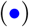 and the Pareto-efficient datasets 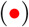. The utopian point is shown by a red star 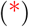. At the utopian point, both goals reach their optimization goal.

Moreover, optimization criteria like the ones considered so far can be grouped to define biologically relevant phenotypes. Grouping criteria into behavioral classes that reflect their compliance with such evolutionary requirements is very helpful to understand the design principles involved. Unbranched pathways, such as our case study, often belong to one of two classes: demand-driven or supply-driven. Pathways producing building blocks such as amino acids are driven by demand, since their metabolic role is to supply as much product as required by cell growth. Overflow metabolism, on the other hand, works as an escape valve for materials that cannot be processed at a certain moment and are therefore required to respond to an excess of precursor. They are therefore supply-driven. Robustness, however, is a requirement for both types of pathway.

#### Profiles of metabolite concentrations

Analysis of the x-ensemble is relatively straightforward in this case due to the simple topology of the pathway. Since the definitions of equilibrium constants are hyperplanes in logarithmic coordinates, the conditions for thermodynamic feasibility define a simplex in design space, a tetrahedron in our case study. Some of the criteria defined above can be ignored for this simple example because they align so well that optimizing one automatically optimizes the others. This is the case for transition time, which is directly proportional to the total concentration of metabolites. Fig. 5 (upper left) shows a criterion space with two conflicting objectives: minimizing total metabolite concentration and the draft of the pathway. They can be simultaneously achieved by minimizing the concentrations of the first two metabolites of the pathway. However, the final product leads to a tradeoff between the two goals, so a Pareto front of optimal solutions arises where *x*_1_ and *x*_2_ are minimal for each possible concentration of *x*_3_. In this particular case, the location of the Pareto front in design space can even be calculated analytically (it is displayed as a red line in the upper right panel of the figure) so no further analysis is required. A further step can be taken by adding another objective, the thermodynamic cost, and defining a Pareto front for all three goals. A two-dimensional projection of this Pareto front is shown in the lower left panel of Fig. 5. The location of this more complete Pareto front in design space is shown in the last panel of the figure and provides a very reduced subset of possible concentration profiles that can be used to build models of our unbranched pathway. In the example shown in the figure, 10,000 concentration profiles where analyzed to obtain a subset of just 140 efficient solutions. Since the operations required for the analysis are extremely simple, an extremely high number of samples can be pre-processed like this before progressing to more demanding tests such as stability analysis. Once a subset of the x-ensemble has been selected, more realistic models can be obtained by formulating all possible combinations of Pareto optimal metabolite concentrations with equally filtered sets of kinetic orders.

**Fig 5.**
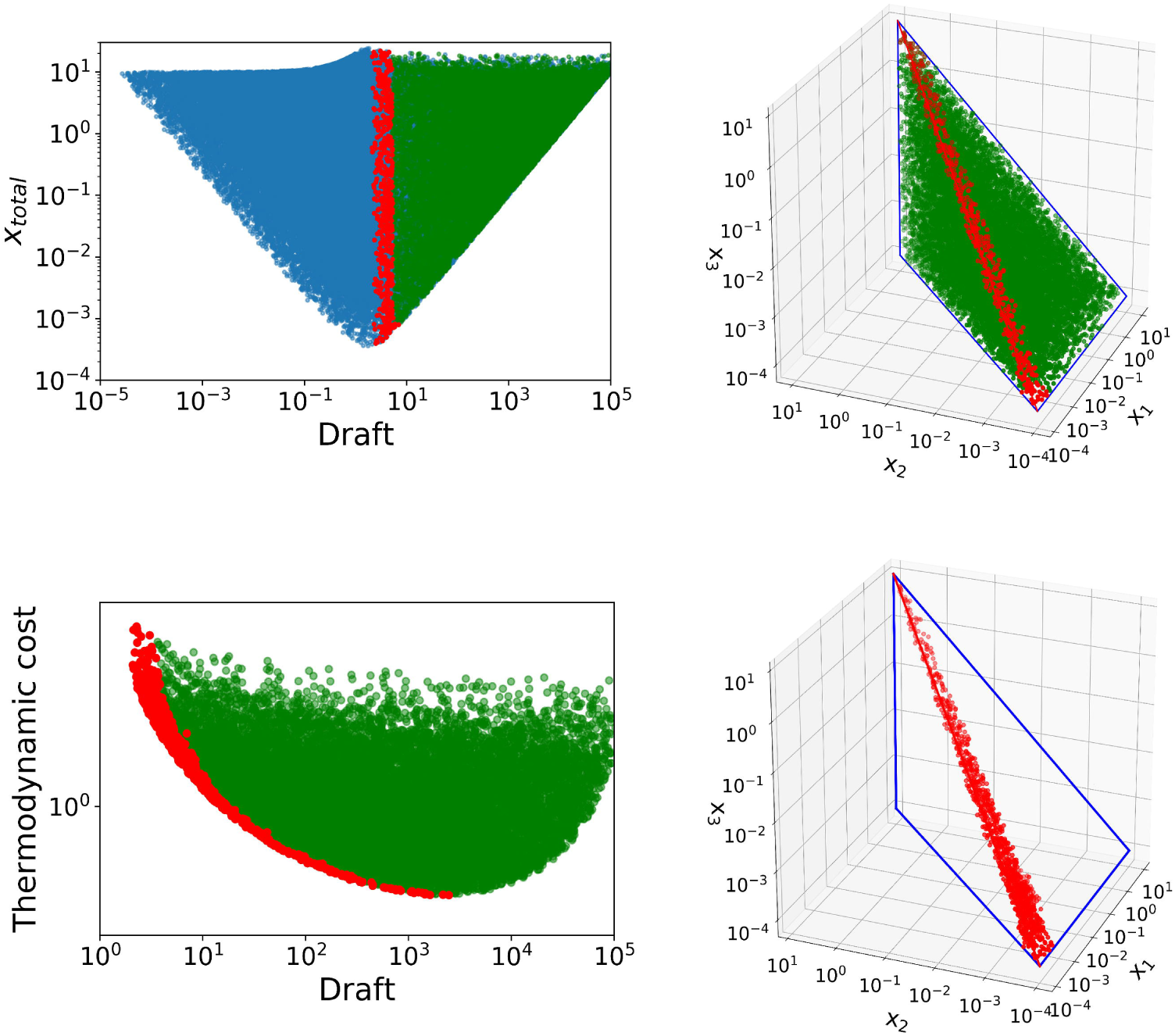
X-ensemble in criterion and design space. The upper left figure shows ensembles in the criterion space with a Pareto front for simultaneous minimization of total metabolite concentration and draft. The upper right panel shows the same data in the design space (unfeasible cases not shown). The lower left panel shows a projection of a different two-dimensional criterion space of draft vs. thermodynamic cost. In this case, the Pareto optimal solutions were calcuated using all three criteria simultaneously. On the lower right panel, only the Pareto optimal solutions are shown in design space. In all design space plots, the feasible area is marked as a tetrahedron and the vertex drawn in red corresponds to the x-total vs. draft Pareto front. Datapoints are colored depending on whether they correspond to thermodynamically unfeasible 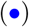, feasible 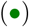 or the Pareto-optimal ensembles 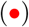.

#### Robust pathways

Once a subset of metabolic concentrations has been selected, kinetic orders can be used to calculate other relevant criteria. Let’s define a robust pathway as the behavioral class that performs above average, hence within the top 50% of the ensemble, for all three robustness criteria: APS, ISS, and ISD. A comparison of the performance of such pathways regarding the other criteria (i.e. the response to supply on the left panel and to demand on the right) is shown in Fig. 6. In regard to supply-driven designs, the robust pathways demonstrate a lower responsiveness, especially concerning the flux. On the demand side, however, there is no conflict regarding robustness. All robust pathways, marked as red points in the figure, have as good a performance as the non-robust cases. Robust pathways have the potential to provide high increases in flux upon demand coupled with low decreases in final product concentration. These results suggest, that there is an inherent conflict between pathway supply orientation and simultaneous robustness.

**Fig 6.**
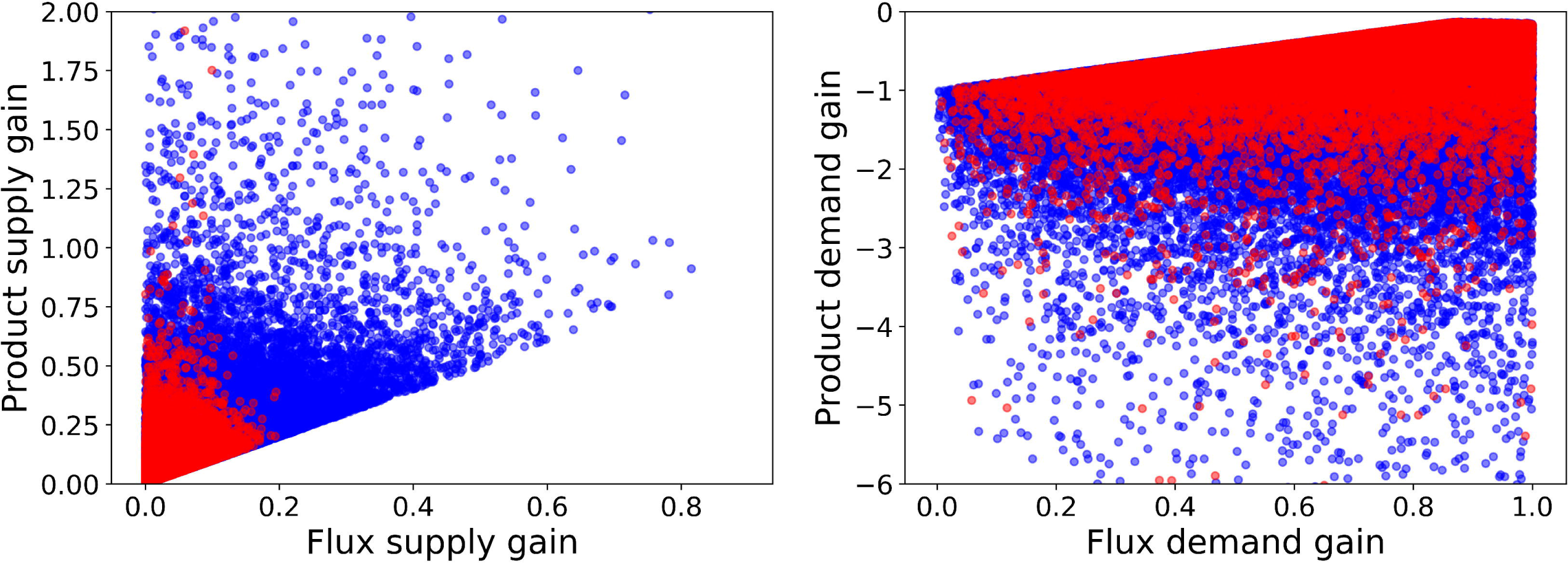
Criterion space projections of supply-driven and demand-driven models. Projections of different models in 2D criterion space related to response to supply (left) and to demand (right). The bulk of models are marked in blue 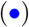, and those models in the intersection of the upper 50% for all three robustness criteria - APS, ISS, and ISD - are superimposed in red 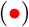.

#### Demand-driven pathways

Since there is no conflict between response to demand and the robust behavioral class, a composite class was defined comprising the above average performers for all three robustness criteria plus the two demand responses (flux and product).

The tradeoff between product and flux response, as depicted in Fig. 7 A, shows that the Pareto front cannot be taken directly as a measure of demand performance. As can be seen in the rightmost edge of the front, marked in red on the figure, negligible gains in flux demand gain are associated with dramatic losses in product demand gain. In this case, the theoretical concept of Pareto optimality is not practical and an alternative must be found. Both criteria were thus combined into a criterion represented in panel B. The new criterion - demand performance - is a linear combination of both original criteria with attributed weights that set the axis of this criterion orthogonal to a fitted line of the Pareto front.

**Fig 7.**
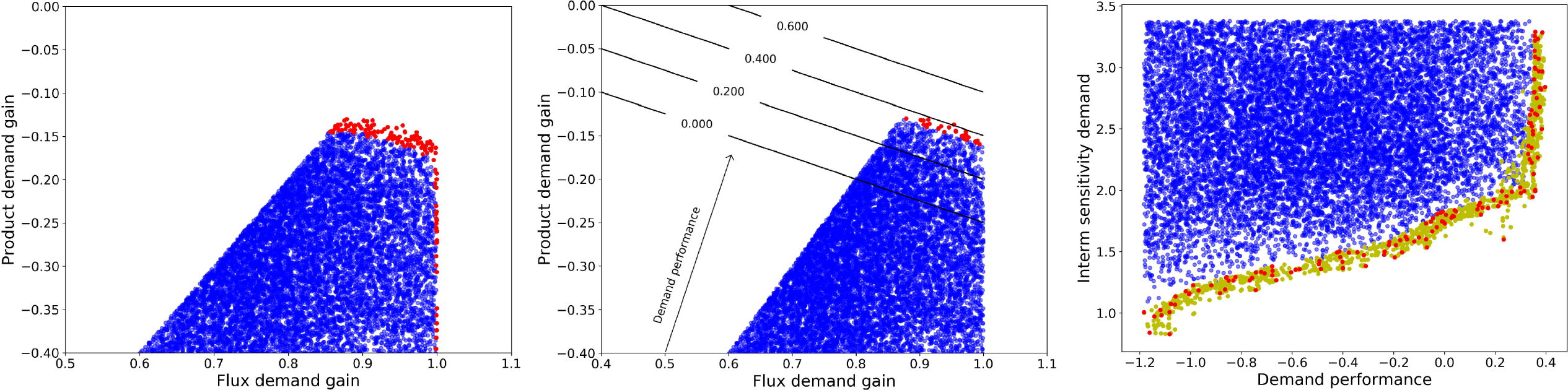
Analysis of demand-driven pathway designs. A: Flux vs. product concentration response to an increase in demand. B: Definition of a new criterion, demand performance, that ensures a balanced response both in flux and in product concentration. C: Pareto front for the tradeoff between demand performance and sensitivity. In all cases, the bulk of the models is represented by blue points 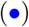, Pareto efficient solutions in red 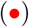, and a Pareto front enriched by convex combinations is shown in yellow 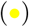.

Within the behavioral class, there is still a tradeoff between the performance of the pathway in terms of demand response and its robustness regarding concentration of intermediates. The corresponding Pareto efficient solutions are marked in red on the third and final panel of Fig. 7 C and a reconstruction of the Pareto front through convex combinations of nearest neighbors is depicted in yellow.

#### Supply-driven pathways

As was shown, defining the robust pathways as a class, robustness and supply response cannot be achieved simultaneously. Testing each robustness criteria separately shows that the source of this conflict is actually the Intermediates Sensitivity to Supply (ISS). For this reason, the behavioral class defined to analyze supply response includes the above average performers for the two non-conflicting robustness criteria: APS and ISD as well as the two criteria related to supply response. Regarding ISS, instead of filtering out all below average performers, which would also eliminate supply responsive designs, only extreme cases (worst 3%) were removed. Further analysis of this class will clearly show the reasons for this conflict.

As shown in Fig. 8, product supply gains can reach very high values in the generated models, especially when the corresponding flux gain is low. Flux supply gain, however, is always below one and any increase in it involves a steep drop in the maximum achievable product gain. The number of samples with high flux supply gains is very low and as the flux supply increases, the range of possible values for the product gain narrows down considerably. This suggests that a delicate balance is needed to increase both flux and product concentration simultaneously. Given that the product gain is actually always positive, and product gains around unity are compatible with any flux gain, maximizing the flux gain is the safest choice to obtain a demand-driven pathway.

**Fig 8.**
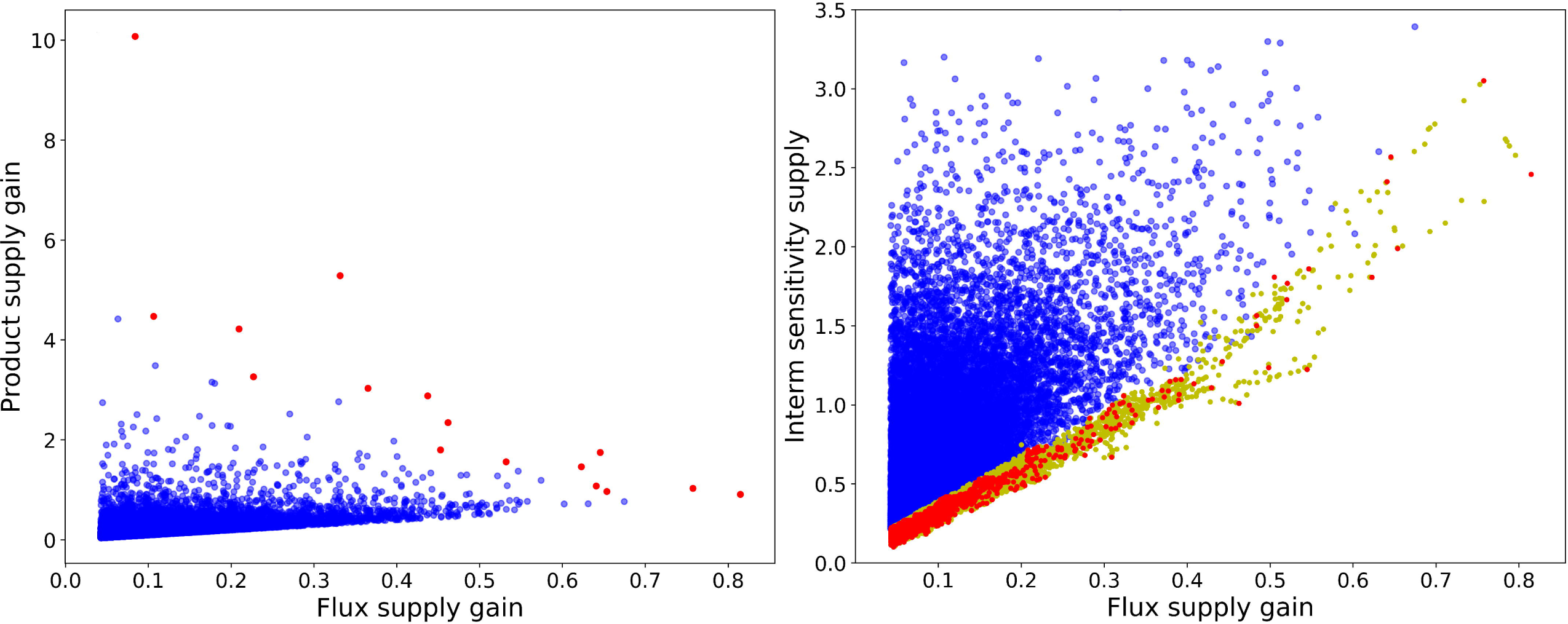
Analysis of supply-driven pathway designs. A: Flux vs. product concentration response of the pathway to an increase in supply. B: Pareto front for the tradeoff between supply performance and sensitivity. In all cases, the bulk of the models is represented by blue points 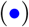, Pareto efficient solutions in red 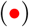, and a Pareto front enriched by convex combinations is shown in yellow 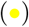.

### Tracking optimal phenotypes back to the design space

Since the initial sampling of kinetic orders was uniform, deviation in their distribution, when considering members of a single class only, provides some insight into how individual parameters influence the behaviour of the system [50]. To further address the question of how to build a robust, yet responsive pathway, we need a method to assess the relative importance of each parameter’s contribution and shed light into potential interactions. Fig. 9 is a very compact display of information relating parameters to their influence on the performance for each criterion. The figure is built by training a logistic regression classifier that attempts to identify which models belong to the class of the top 50% performers for each criterion individually. Fig. 9 shows the vectors defining the separating hyperplane for each criterion. Parameters and criteria are ordered in the cladogram according to the similarity of their coefficient profiles. Parameters (criteria) clustered together show similar influences (or responses).

**Fig 9.**
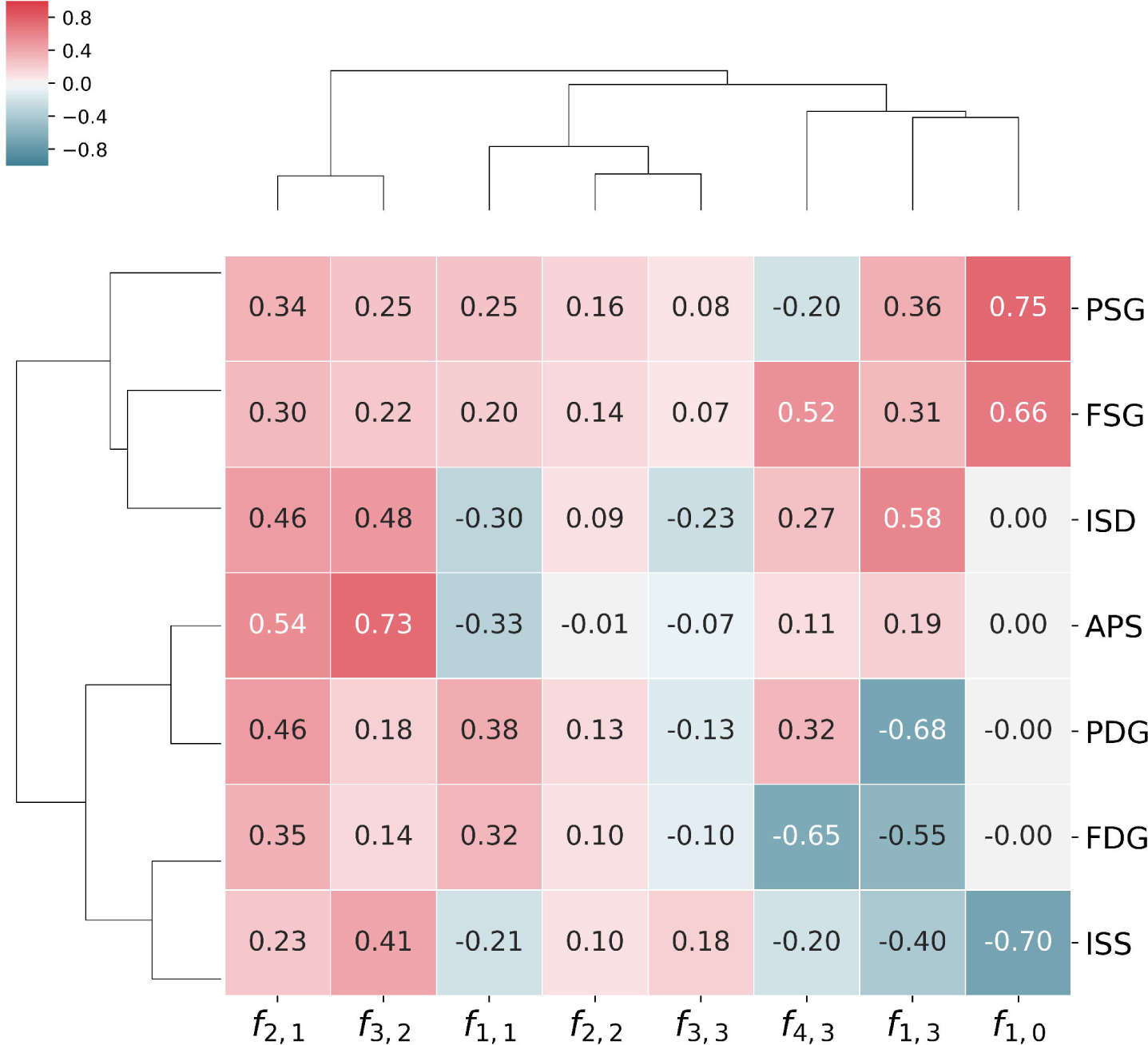
Parameter values of the logistic regression model. The influence of each β-value (model parameter) on the respective goal and kinetic order is shown. Coefficients for logistic regressions to classify the the top 50th percentile for each analyzed goal: Intermediate sensitivity to supply (ISS), intermediate sensitivity to demand (ISD), aggregate parameter sensitivity (APS), product supply gain (PSG), flux supply gain (FSG), product demand gain (PDG) and flux demand gain (FDG).

In accordance with what is known about unbranched pathways, kinetic orders of the backwards reactions *f*_2,2_ and *f*_3,3_ are generally of little importance, as is reflected by the smaller magnitude of their coefficients in the table. An exception to this rule is the kinetic order of the first reaction for its product, *f*_1,1_. Its coefficients are larger in magnitude reflecting this parameter to be important while the negative signs for all three sensitivity criteria show how an irreversible first step (lower sensitivity to its own product) results in more robust designs. Large kinetic orders for the forward reactions *f*_2,1_ and *f*_3,2_ tend to improve all goals and are often significant. Finally, the conflict between intermediate sensitivity and supply response can also be clearly seen in the figure. The profiles for APS and ISD are very similar to one another and to those for both supply gains – PSG and FSG – but that of ISS is clearly different, being more similar to those for demand gains – PDG and FDG. It is therefore intuitive that ISS will be better aligned with demand-oriented goals and lead to conflicts with supply-oriented goals. Moreover, the profile for ISS is dominated by three parameters: *f*_3,2_, *f*_1,0_ and *f*_1,3_. The first parameter, *f*_3,2_, has positive coefficients for every goal, so it unlikely to be the cause of any conflict. A completely different situation can be appreciated regarding the the saturation of the first reaction *f*_1,0_ by the precursor, which is also the dominant parameter for the supply gains PSG and FSG, albeit with coefficients of opposite signs. Finally, the strength of the feedback loop *f*_1,3_ has high magnitude coefficients for almost all criteria and opposing signs for different goals. A divide would be expected here with strong feedback loops (more negative values) being a characteristic of pathways robust to supply with better demand gains, PDG and FDG, while weaker feedback loops (values closer to zero) would be characteristic of pathways robust to changes in demand and more responsive to supply. This is already a clear indicator that end-product inhibition is rather a design principle for demand-driven pathways. Supply-driven pathways must rely on different mechanisms. The saturation of the final step, also called elasticity of demand, does not play a major role for any of the robustness measures, but it is important for tradeoffs between the flux responses to supply and demand.

The approaches presented so far help identify the key parameters and relate their values to the performance on different criteria but identifying a design goes beyond that. Given a region on the criteria space, one should be able to identify a region in design space able to meet those specifications. This area can be directly expressed as one or more constraints on the original parameters, but a more compact description can be achieved by defining a new set of coordinates based on Principal Component Analysis (PCA) of the models in the selected region. PCA provides a transformation that groups together those variables that are linearly correlated in the selected dataset and orders them according to variance. The groups of parameters that vary the most in the selected set will be grouped in the first few Principal Components (PCs) while those that remain approximately constant will be grouped in the last few. Finally, the new variables will be orthogonal to one another making them specially suited for independent sampling.

As an example, we will use PCA to try and map the Pareto front in Fig. 7 to a region in the design space. First, PCA is applied to the enriched Pareto front. The new coordinates will still span the full design space but will be centered at the mean of the parameter values that can be found in the Pareto front and grouped by correlation. Now the Pareto front is divided in two approximately convex sets (the two roughly linear segments in Fig 10) and regression is used to find correlations between the PCs within those sets. The first segment shows strong correlation between the *PC*_1_ and *PC*_2_ while the second shows a strong dependence between *PC*_1_,*PC*_4_ and *PC*_7_. Thus, a parametric equation can be generated to approximate the first segment by varying *PC*_1_ as a free parameter and adjusting *PC*_2_ accordingly. The same can be done for the second segment with two parametric equations, one for *PC*_4_ and *PC*_7_. Fig 10 shows such segments.

**Fig 10.**
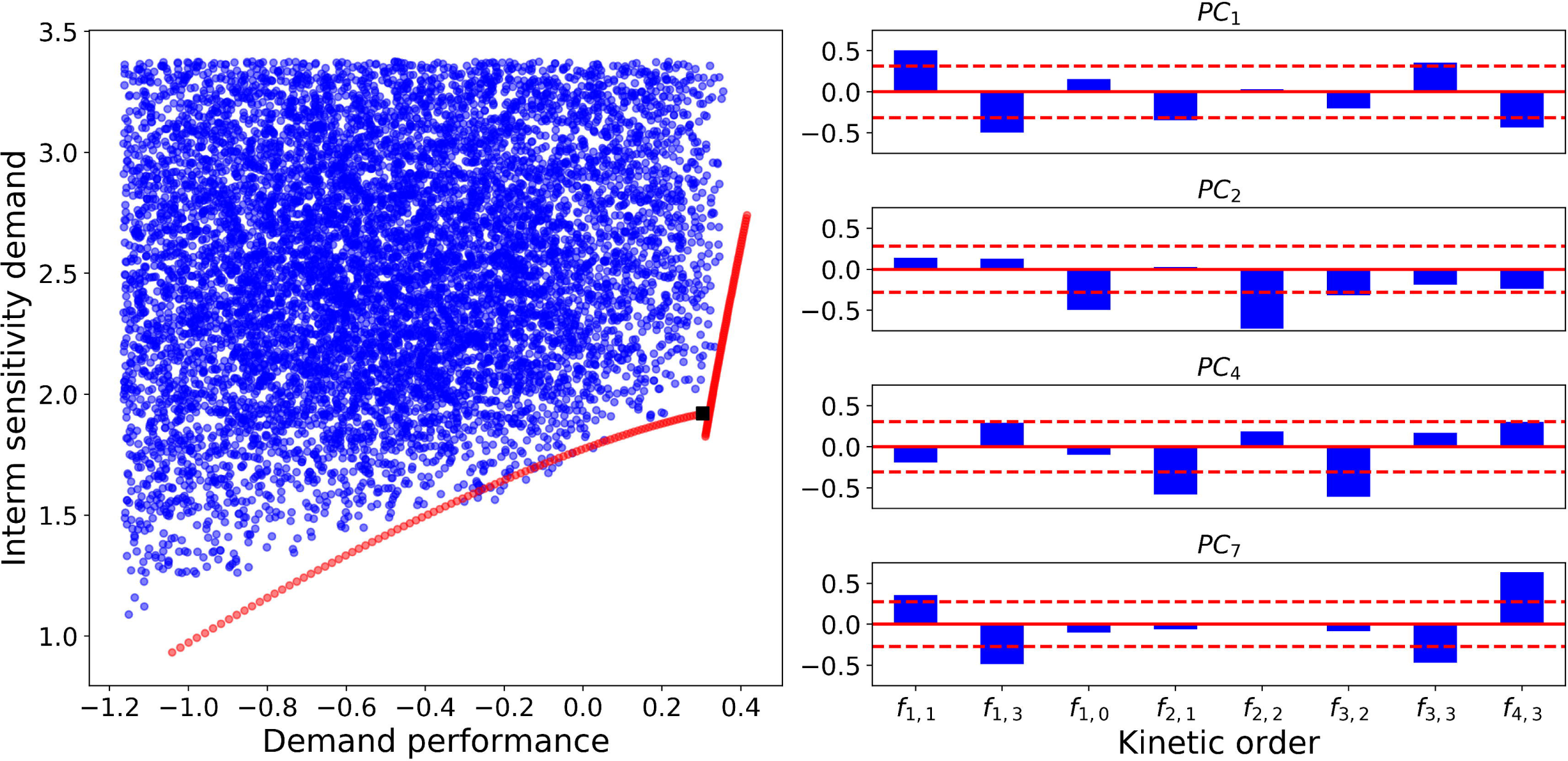
Datasets synthesized by PCs. **Left:** The synthesized data are shown in the criterion space. The location of designs generated by direct sampling in PCA coordinates 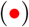 is compared with non-efficient datasets 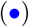. The synthetic data were generated as two independent segments. In both cases, the first principal component was varied as a free parameter, while a minimal set of other PCs were varying according to how they correlated in the corresponding segment of the Pareto front. For the first segment (horizontal trend), PC2 that was varied linearly to the function *PC*_2_ = −0.70*PC*_1_ − 0.83, while all other PCs were kept constant. For the second segment, two PCs other than PC1 (PC4 and PC7) had to be varied according to the following fucntions: *PC*_4_ = 0.95*PC*_1_ − 2.07 and *PC*_7_ = −0.54*PC*_1_ + 1.18. Thus, the segments were obtained through the variation of only two and three PCs respectively. **Right:** The right panel shows the relevant principal components as bar plots of the coefficients with which each kinetic order contributes to the corresponding PC. The dashed red lines show the average value of all the coefficients in absolute value. Thus, the relevant kinetic orders for each PC are identified as those with coefficients exceeding the dashed lines, such as *f*_2,2_ in *PC*_2_.

### Stability analysis and dynamic simulations

Once the x- and the k-ensemble have been screened separately using computationally less intensive tests, all combinations between both ensembles can be generated to test for stability. In this case, a selection of 140 cases from the x-ensemble was combined with 1,060 cases selected from the k-ensemble as demand-driven pathways and 20,360 selected as supply-driven pathways. The two complete ensembles contained 148,400 models of demand-driven pathways and 2,850,400 supply-driven. The fraction of unstable models was well below 1% in all cases, its number increasing with pathway length and feedback intensity as has been described elsewhere [4]. Moreover, saturation of reactions by their products (kinetic orders close to zero) also seems to contribute to instability (data not shown). Once the unstable cases have been discarded, a full set of models is available that has already been classified for a series of phenotypic traits and can further be analyzed through other means. Since the topic of stability analysis of ensembles of models has already received some attention and any of the known approaches can be easily integrated in our pipeline, we will refer the reader to the available literature on the topic [12, 51]. It is worth noting, however, that generalizing the methods outlined above to recognize features linked to stability in the joint v-x-k-ensemble would be straightforward.

Numerical integration for some models selected from different regions of the criterion space shows the type of behavior anticipated by selection of the k-ensembles. Models 1, 2 and 3 in Fig. 11 have increasingly better responses to an increase in demand, but they also have an increasing tendency to accumulate intermediates. Model 4 performs poorly for all criteria, since it has been taken from a region far away from the Pareto front.

**Fig 11.**
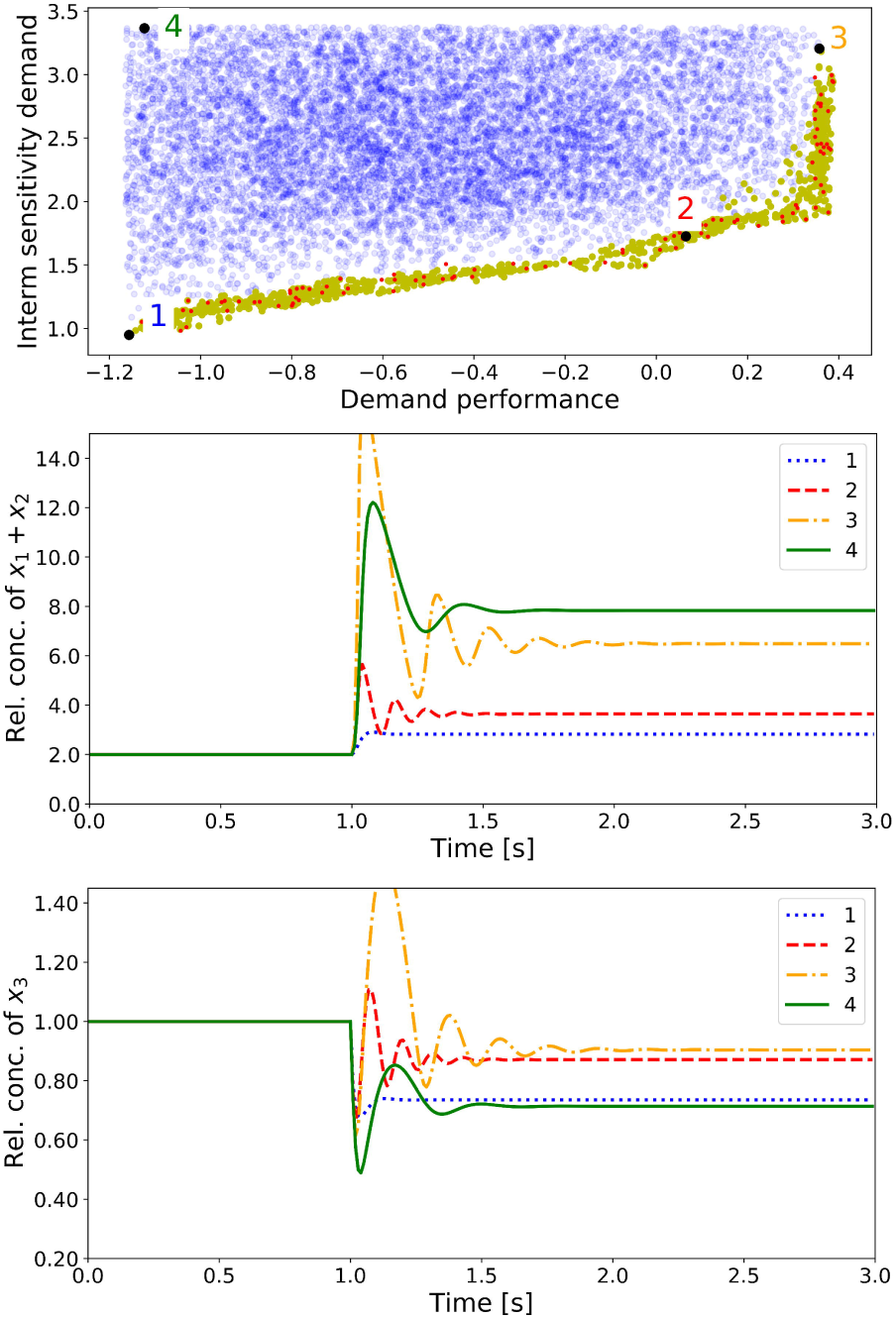
Dynamic simulation of perturbed pathway designs. Models of demand-driven pathways are chosen and visualized in the criterion space with the respective Pareto front of ISD vs. demand performance. Simulations are performed using the normalized system with a reference state, setting/fixating the initial steady state concentration at 1. After one second, the demand factor *x*_4_ is increased by 100%. Three (1-3) designs are chosen from distinct regions along the Pareto front and one (4) was selected from a region away from the front. The dynamic response of the different models is visualized in for the sum of the intermediate concentrations *x*_1_ + *x*_2_ (middle panel) and for the product concentration *x*_3_ (bottom panel).

## Discussion

Formulating and understanding dynamic models is an important prerequisite for systems and synthetic biology. The current trend of integrating information from many different sources to obtain such models has led to important advances [15] but the challenge of working under high uncertainty remains.

We have shown that much is to be gained by replacing the traditional kinetic parameters such as saturation constants and *V*_*max*_ by a standardized set based on easier to measure magnitudes (steady state fluxes and concentrations) and phenomenological kinetic coefficients (kinetic orders a.k.a. elasticities). This choice of parameters enables a seamless integration of kinetic analysis with flux balancing and thermodynamics but it also has the additional advantage of providing a better set of defining features to apply pattern searching algorithms. This is so because the global response of a system is a more direct function of the kinetic order than of each individual kinetic parameters. But the greatest benefit that results from this parameter choice is, by far, the possibility of screening phenotypes using only a subset of the full parameter space. Partitioning the design space in several ensembles has an opposite effect to the combinatorial explosion that is often called the curse of dimensionality, not countering it completely but at least mitigating it. Finally, this choice of parameters does not result in any loss of generality since a model thus defined can always be translated to more traditional kinetics.

Pareto optimality provides insight on how biological systems can find tradeoffs between conflicting goals. This approach has been extremely successful in cases where the mapping between design and criterion space fulfills certain conditions. In many cases, the performance of a certain phenotype for each criterion is a monotonic function of its distance to the best performer for such goal [38, 39, 52]. In such cases, the image of the Pareto front in design space is a convex combination of a handful of specialized phenotypes or archetypes. Each archetype is the best performer for a given criterion considered alone. For the general case of biochemical networks, the mapping between design and criterion space is not normally monotonic. In fact, it is is not even continuous, thus hampering a direct reconstruction of the Pareto front in the design space. The methods presented here intend to be one more step towards an understanding of biochemical networks on the basis of biological first principles.

Although our emphasis is on finding general design principles, this method can be prepended to any modeling pipeline as a way to accelerate parameter estimation by filtering our bad models and integrating qualitative biochemical knowledge into the modeling process. In fact, the method presented here can be considered the natural continuation of the ensemble modeling approaches developed by Liao’s group [13, 14] by way of incorporating the attention to thermodynamics and the use of power-law kinetics exemplified by Hatzimanikatis’ lab [15, 21, 29]. What this method adds is the possibility of partitioning the design space into subspaces that can be sampled and tested independently. This possibility expands the applicability of ensemble approaches to much larger metabolic networks. In this respect, we are proposing a complementary approach to that of Design Space Analysis put forward by Savageau and coworkers [24, 25]. This latter approach achieves a much deeper knowledge of the system without the need for sampling by dividing the design space into qualitatively similar subspaces. Since the number of possible subspaces grows exponentially with the size of the network, the possibility of developing hybrid approaches that combine the scalability of our method and the detailed results of DSA is one worth looking into.

The particular example chosen to test the method is by necessity very simple, yet all its properties derive from extremely non-linear interactions. Nevertheless, the analysis shown here performs extremely well using only linear methods (classifiers based on hyperplane separation and PCA) and it deserves further comment. We have intentionally applied the simplest possible tools from machine learning to ensure a robust identification of the most important trends in a manner that can be understood in biological terms. To apply this method with emphasis on prediction rather than on description, more sophisticated methods can be used. A simple logarithmic transformation of the parameters and the incorporation of interaction terms with regularization results in a much more accurate classification of classes at the expense of more complex relationships between the parameters. Exploring this tradeoff is definitely a worthwhile task ahead of us.

Finally, stability becomes a much more relevant issue when ensemble approaches are applied to larger models. In this case, the fraction of the design space that leads to stable systems is normally smaller and identifying stable regions becomes a priority. The process of selection based on partial ensembles – v,x and k – can alter the fraction of stable systems once all the ensembles are recombined to the final v-x-k-ensemble of models that can be tested for stability. If the resulting ensemble has a higher fraction of stable states than a randomly generated one, not much remains to be done except further filtering using the methods already discussed and those previously available [12, 51]. In case the pre-selection decreases the fraction of stable models, further trade-offs can be found and the goals conflicting with the stability of the system can be detected. The partitioning of parameters into subspaces can also pinpoint the causes of instability. For instance, saturation of certain enzymes has been identified as relevant for stability in this and other studies [22, 51], but separating the effect of saturation and other destabilizing interactions between kinetic parameters and concentration profiles can be extremely difficult in ensembles based on traditional kinetic parameters such as saturation constants. The partitioning proposed here reflects saturation exclusively as a property of the k-ensemble. Since the Jacobian of the system is defined by the kinetic orders and the turnover numbers, it is extremely easy to establish whether unstable systems correlate with saturated enzymes - kinetic orders close to zero - or they are rather linked to the timescales of the system represented by the turnover numbers [53]. So, even though stability analysis can only be performed on the full ensemble, the previous analysis of its component subspaces can provide a great deal of insight.

## Conclusion

The concept of design principle has been part of the biological lexicon [45] as long as its counterpart, design pattern, has been in architecture, [54] and much longer than in software engineering [55]. The concept has, however, not been as influential in biochemistry as it has been in the above mentioned disciplines. This can in part be explained by the initial reluctance of the biological community to accept quantitative methods. After all, it took almost thirty years for Systems Biology to become part of the mainstream [56]. But as Systems and Synthetic Biology mature, it would be expected that design philosophies involving design principles / patterns will emerge. Following in the footsteps of other disciplines, we could expect design principles to be catalogued in a similarly structured manner – see [54, 55] – with each principle being presented with a clear name, a description of the problem it solves, details of how to implement it, and the tradeoffs that it causes as well as possible drawbacks to be considered. Moreover, design principles are meant to be used together, complementing one another. In architecture, for instance, patterns have a hierarchical structure that goes from organization at a high level, such as **distribution of towns** in a region, all the way down to organization of a house in patterns like **sleeping to the east** or **windows that open wide**.

In the case of biochemical networks, a good amount of design principles have already been catalogued [45, 57, 58] so we can expect a higher degree of formalization and structure in the future. In our case study, we can see how known patterns naturally emerge together and shape unbranched pathways. A **supply-** or **demand-driven pathway** can satisfy different metabolic needs. We have shown how **end-product inhibition** can help implement a demand-driven pathway, but hamper the design in the supply-driven case. A detailed implementation of the unbranched pathway with end-product inhibition can involve a **proportional feedback** acting on an **irreversible first step** that is far from equilibrium to guarantee the efficiency of the feedback [44], but also one that is saturated by its product so that the corresponding kinetic order is close to zero [43]. The pathway benefits from all other steps having non-zero kinetic orders for both their substrates and their products, which can be achieved by a design principle that is well known by biochemists and could be named **S near Km** [59]. Regarding tradeoffs and drawbacks, increasing the intensity of the feedback improves the performance of the pathway at the cost of reducing its stability margin and this gets worse as the length of the pathway increases [43, 45]. The pathway can be kept within the stability margin by the application of two further design principles: **kinetic shortening** [45] and **thermodynamic shortening** [31]. Regular formalisms combined with ensemble modelling can take advantage of the recent developments in machine learning, thus helping to find more design principles as well as to understand how they come together in complex biochemical networks. For instance, all the design principles mentioned above have been identified, one at a time, during the last few decades. The analysis of this simple pathway, using the method proposed here, not only shows all these principles emerging together from the analysis Fig. 9, but also shows clearly how they are linked to demand driven pathways and would never work properly to obtain a supply driven system Fig. 6. This hierarchical grouping of known design principles and their subordination to a demand driven role has not, to our knowledge, been observed before.

## Supporting information

**S1 Appendix Detailed mathematical model and notes on sensitivity analysis**

## Acknowledgments

A.M.S. was funded by the German Ministry of Education and Research (BMBF) project HOBBIT (031B0363A).

